# Genetically-modified *X. laevis* indicate distinct roles for prominin-1 and photoreceptor cadherin in outer segment morphogenesis and retinal dystrophy

**DOI:** 10.1101/2020.08.23.263780

**Authors:** Brittany J Carr, Paloma Stanar, Orson L Moritz

## Abstract

Mutations in prominin-1 (*prom1*) and photoreceptor cadherin (*cdhr1*) are associated with inherited retinal degenerative disorders such as retinitis pigmentosa, cone-rod dystrophy, and juvenile macular dystrophy. The proteins encoded by these genes are hypothesized to regulate photoreceptor outer segment disc morphogenesis, but their functions remain unknown. We used CRISPR/Cas9 to generate *prom1-, cdhr1-*, and *prom1* + *cdhr1*-null *X. laevis* and then documented the effects of these mutations on photoreceptor structure and function. *Prom1*-null mutations resulted in dysmorphic photoreceptors comprised of overgrown and disorganized disc membranes. Cones were more severely affected than rods; outer segments were elongated and fragmented, and ERG response was impaired. Autofluorescent deposits in the outer segment layer of aging *prom1*-null animals indicate that secondary toxic effects to the retina or RPE drive retinal degeneration for this mutation, instead of direct effects on outer segment disc morphogenesis. *Cdhr1*-null photoreceptors did not appear grossly dysmorphic, but ultrastructural analysis revealed that some disc membranes were overgrown or aligned vertically within the plasma membrane. *Prom1* + *cdhr1*-null mutants did not differ significantly from *prom1*-null mutants. Our results indicate that neither prom1 nor cdhr1 are necessary for outer segment disc membrane evagination or the membrane fusion event involved in disc sealing. Rather, they are necessary for higher-order organization of the nascent outer segment discs. Prom1 may align and reinforce interactions between the disc leading edges, a function more critical in cone photoreceptors for structural support. Cdhr1 may help to align nascent discs and maintain horizontal disc orientation prior to fusion.

## INTRODUCTION

The retina is a light-sensitive neural tissue that initiates vision though rod and cone photoreceptors, which are named for the shape of their outer segments (OS) – a highly-modified primary cilia. Rod outer segments (ROS) are cylindrical and comprised of stacks of discrete membranous discs sheathed in a plasma membrane. Cone outer segments (COS) are tapered and comprised of open discs formed by infoldings of a continuous membrane. OS morphogenesis is dynamic; new membrane discs are synthesized continuously by plasma membrane evaginations at the basal ROS and COS (1–4). Deficiencies in this process can result in retinal degenerative disease (5).

Inherited retinal degenerative diseases are rare disorders that result in progressive vision loss and blindness. Defects in prominin-1 (*prom1*) and cadherin-related family member-1 (*cdhr1*) can result in autosomal dominant Stargardt-like macular dystrophy (6, 7) or autosomal recessive cone-rod dystrophies and/or retinitis pigmentosa (8, 9). The proteins encoded by these genes share a unique localization to the basal ROS and COS where nascent discs are synthesized, which has lead to the hypothesis that mutant *prom1* and *cdhr1* cause retinal degeneration by disruption of OS morphogenesis (Rattner et al., 2001; Zacchigna et al., 2009; Han et al., 2012).

*Prom1* – a.k.a. prominin-like 1, CD133, or AC133 – encodes a pentaspan transmembrane glycoprotein that is evolutionarily conserved between vertebrates and invertebrates (12). It is hypothesized to be involved in ectosome formation (13), neurodevelopmental signalling (14), stabilization of membrane curvature (15), and cytoskeletal remodelling (16). Interestingly, however, the only documented clinical manifestation of *prom1* mutations is non-syndromic vision loss (9). *Prom1*-null mice exhibit dysmorphic ROS and COS, mislocalized rod and cone opsins, increased photoreceptor apoptosis, and impaired retinal function; retinal degeneration occurs within 3-26 weeks, depending on the genetic background of the mouse model used (11, 17). ROS disorganization, lipofuscin-like deposits in the RPE, a reduced scotopic and photopic electroretinogram B wave, and retinal degeneration also occur, by 4-7 weeks, in mice with overexpression of the autosomal dominant mutation, *prom1^R373C^* (18). It has been suggested that prom1 and cdhr1 proteins form a complex that performs a shared role in OS morphogenesis because cdhr1 is mislocalized in *prom1^R373C^* mice, mouse prom1 is mislocalized in *cdhr1^-/-^* mice, and prom1 coimmunoprecipitates with cdhr1 in HEK293 cells cotransfected with either wildtype or mutant prom1 and a cdhr1-Myc fusion construct (18).

C*dhr1* – cadherin-related family member 1 a.k.a protocadherin-21 (*pcdhr-21*) or photoreceptor cadherin (*prCAD*) – encodes a single-pass transmembrane protein expressed exclusively in the retina; orthologues have been identified in mammals, birds, fish and amphibians, but not invertebrates (8). *Cdhr1*-null mice have dysmorphic ROS and increased retinal apoptotic activity and degeneration by 6 months (8). Mammalian cdhr1 is proteolytically-cleaved in the photoreceptors, resulting in an extracellular soluble N-terminal fragment and a C-terminal fragment that remains embedded in the plasma membrane. It has been hypothesized that cleavage of cdhr1 may drive OS assembly by acting as an essential, irreversible step, such as the “release” mechanism during plasma membrane fission in ROS that seals off discs (19). Alternatively, cdhr1 has been reported to act as a tether between the leading edge of nascent ROS discs and the inner segment (2).

Despite studies demonstrating that loss of *prom1* or *cdhr1* causes photoreceptor OS defects and retinal degeneration in mice, their roles in OS morphogenesis and the mechanisms of pathogenicity remain unknown. Here, we report the characterization of genetically-modified *prom1*-null, *cdhr1*-null, and *prom1 + cdhr1-null X. laevis*, which provide new insights into the roles of these proteins in photoreceptor OS morphogenesis and retinal degenerative disease.

## METHODS

### Animal ethics statement & housing

Animal use protocols were approved by the University of British Columbia Animal Care Committee and carried out in accordance with the Canadian Council on Animal Care and the ARVO Statement for the Use of Animals in Ophthalmic and Vision Research. Animals were housed at 18°C under a 12-hour cyclic light schedule (900-1200 lux).

### RNA identification, construction, and synthesis

Single-guide RNAs (sgRNAs) were synthesized on the basis of *X. laevis prom1* and *cdhr1* sequences identified by xenbase.org (*prom1*: xelaev18005149m, NCBI Gene ID: 100316924 (*proml-1*); *cdhr1*: xelaev18035010mg, XB-GENE-865231, NCBI Gene ID: 100337587 (*cdhr1.L*) & xelaev18000599mg, XB-GENE-17339736, NCBI Gene ID: 108703385 (*cdhr1.S*). SgRNAs, corresponding oligonucleotides, and primer sequences were designed *in silico* using ChopChop (http://chopchop.cbu.uib.no/), ZiFiT (http://zifit.partners.org), and Integrated DNA Technologies OligoAnalyzer (idtdna.com/calc/analyzer) online tools. The sgRNA target sequences, corresponding complementary oligonucleotides for cloning into the pDR274 vector, and PCR primer sequences used for subsequent genotyping are listed in Table 1.

**Table 1.**
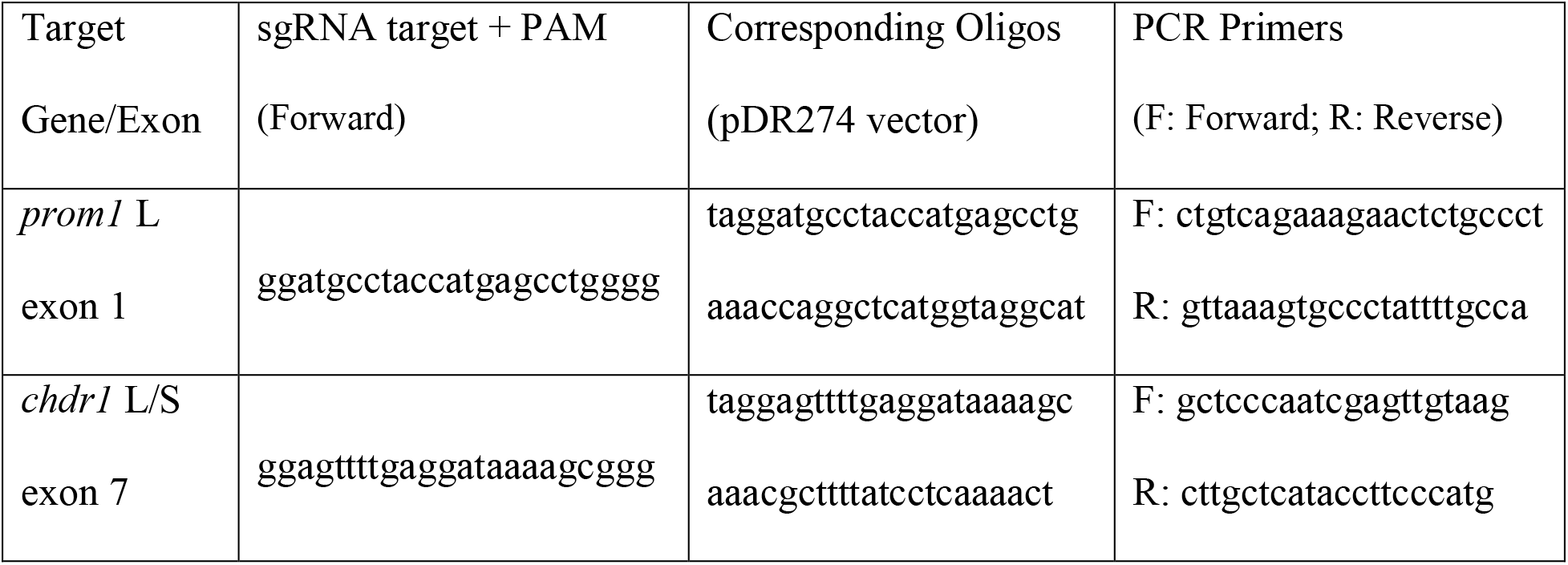
SgRNA targets, corresponding complementary oligonucleotides, and genotyping PCR primers used in this study.

To synthesize sgRNAs, oligonucleotides encoding targeting sequences were cloned into pDR274 (a gift from Keith Joung - Addgene plasmid #42250) and the resulting derivatives were linearized and used as templates for *in vitro* RNA transcription with the HiScribe™ T7 High Yield RNA Synthesis Kit (NEB, Ipswich, MA, USA); 1.5 μg template was incubated for 4 hrs following kit protocols. Cas9 mRNA was transcribed *in vitro* from linearized pMLM3613 (a gift from Keith Joung - Addgene plasmid #42251) using the HiScribe T7 ARCA mRNA kit with poly-A tailing (NEB, Ipswich, MA, USA). A combination of 3 separate reactions was used to achieve high concentrations (≥ 45 μg) of Cas9 mRNA. eGFP mRNA was transcribed *in vitro* from a linearized pBluescript II SK+ construct using the T7 mMessage Ultra kit (Ambion/ThermoFisher Scientific, Waltham, MA, USA). All synthesized RNA was treated with DNase I (NEB, Ipswich, MA, USA) to remove template contamination, and then purified using the Qiagen RNeasy kit (Hilden, Germany). Final products were quantified by absorbance at 260/280 nm with a NanoDrop 2000c spectrophotometer (ThermoFisher Scientific, Waltham, MA, USA), evaluated for size and quality by agarose gel electrophoresis (1.5%), and stored at −80°C prior to use.

### Microinjections, embryo selection, and tadpole rearing

*In vitro* fertilization and microinjections were performed at 18°C as previously described (20). Eggs and sperm were incubated in a petri dish for 20 min, the follicle cell sheath was removed from fertilized embryos using 2% cysteine and gentle shaking, and then the embryos were tightly packed into a monolayer in 2% agarose injection plates flooded with 0.4x MMR + 6% Ficoll. Cas9 mRNA (5 ng), eGFP mRNA (750 pg), and sgRNAs (1.25-2 ng) were combined and loaded into a pulled glass micropipette with a 20–25 μm bore. Micropipettes were mounted in a micromanipulator and connected to 25 μL Hamilton gastight syringes mounted in a Hamilton syringe pump set to deliver 36 μL/hour. Embryos were injected with the RNA solution for 1s (equal to ~10 nL). Injected embryos that exhibited symmetrical division at the 4-cell stage were transferred to 6% Ficoll + 10μg/mL gentamycin in 0.1x MMR at 18°C. At 1-3 dpf, surviving embryos were screened for eGFP fluorescence using an epifluorescence-equipped Leica MZ16F dissecting microscope; uniformly fluorescent embryos were selected and transferred to 1x tadpole ringer (10 mM NaCl, 0.15 mM KCl, 0.24 mM CaCl_2_•H_2_O, 0.1 mM MgCl_2_•H_2_O). Tadpoles were raised in a 12:12 cyclic light incubator at 18°C.

### Confirmation and characterization of CRISPR-mediated indels

Whole embryos (1-3 dpf) or tail snips (14 dpf and older) were used for genomic DNA extraction. Tissues were placed in 75 μL of genomic prep buffer (50 μM Tris-HCl pH 8.5 + 1 μM EDTA + 0.5% Tween 20 + 0.2 mg/mL proteinase K) and incubated at 55°C for 2 hrs and 95°C for 10 min. Extracted genomic DNA was used as template for PCR amplification of target exon sequences using the primers listed in Table 1. Primers and free nucleotides were removed by ExoSAP enzyme treatment (37°C for 30 min and then 95°C for 5 min) and the products were analyzed by Sanger sequencing (Genewiz, Seattle, WA).

### Fluorescence immunohistochemistry & Lucifer yellow dye

For fluorescence immunohistochemistry, whole eyes were fixed in 4% paraformaldehyde + 3% sucrose in 0.1 M sodium phosphate buffer (PB; pH 7.4) overnight at 4°C. Fixed eyes were then cryoprotected in 0.1 M PB + 20% sucrose (pH 7.4) for 3 hrs at 22°C with gentle shaking or 18-48 hrs at 4°C without shaking. Cryoprotected eyes were embedded in Optimal Cutting Temperature medium (OCT; ThermoFisher, Waltham, MA, USA) and then quick-frozen to −80°C. Sagittal or coronal cryosections of the central retina were cut at 12 μm, thaw-mounted onto Fisherbrand Superfrost Plus slides and stored at −20°C until use.

For immunolabelling, sections were washed (3 x 8 min) in 1x phosphate buffered saline (PBS) and then incubated in blocking buffer (1% goat serum + 0.1% TX-100 in 1x PBS) for 30-45 min. After blocking, sections were washed and then incubated overnight in primary antibody in dilution buffer (0.1% goat serum + 0.1% TX-100 in 1x PBS). After incubation, sections were washed and then secondary antibodies and counterstains were applied in dilution buffer and incubated for 4-6 hrs at 22°C; all tissues were counterstained with Hoechst 33342 and fluorophore-conjugated wheat germ agglutinin (WGA) to visualize nuclei and photoreceptor OS membranes. Sections were cover-slipped using Mowiol mounting medium (Millipore Sigma, St. Louis, MO, USA) and imaged using a Zeiss 510 or 800 confocal microscope equipped with a 40x N.A. 1.2 water immersion objective or a Zeiss LSM 880 with Airyscan equipped with a 63x 1.4 N.A. oil immersion objective. Sections were double-labelled in pairs – rhodopsin/cone opsin, cdhr1/prph-2, prom1/WGA – to maximize tissue use from small tadpole eyes. Antibody sources, counterstains, and concentrations used are listed in Table 2. Micrographs represent maximum intensity projections of whole retinal sections (z = 0.28 μm/step) unless specified otherwise.

**Table 2.** Antigens, species/type, source, and working dilutions of antibodies used in this study.

| Antigen | Species/Type | Source | Dilution |
| --- | --- | --- | --- |
| Primary Antibodies |  |  |  |
| *X. laevis* prominin-1 (N-terminus) | rabbit polyclonal | D.S. Papermaster | IHC: 1:750<br>WB: 1:1000 |
| *X. laevis* cone opsin (red-sensitive) | rabbit polyclonal | O.L. Moritz | IHC: 1:500<br>WB: 1:1000 |
| *X. laevis* peripherin-2 (2A5) | mouse monoclonal | R. S. Molday | IHC: 1:10 |
| *X. laevis* rhodopsin (B630N) | mouse monoclonal | P. Hargrave | IHC: 1:15<br>WB: 1:15 |
| *X. laevis* photoreceptor cadherin (N-terminus) | rabbit polyclonal | O.L. Moritz | IHC: 1:10,000<br>WB: 1:10,000 |
| Secondary Antibodies |  |  |  |
| conjugated WGA | AF488, AF594, AF647 | Life Technologies | 1:100 - 1:1,000 |
| Hoechst 33342 | nuclear stain | Sigma Aldrich | 0.1 mg/mL |
| AF488 | anti-mouse<br>anti-rabbit | Jackson<br>ImmunoResearch | 1:750 |
| Cy3 | anti-mouse<br>anti-rabbit | Jackson<br>ImmunoResearch | 1:750 |

To label photoreceptor disc membranes with Lucifer yellow, a “peeled grape preparation” of whole eyes was performed. This is suitable for tadpoles and froglets, but works best in animals whose scleras have not been hardened by cartilaginous growth. Using a 30 gauge needle and extra fine forceps, the sclera was split open and then gently peeled away, along with the RPE, leaving a globe comprised of the retina and intact lens. The retina-lens globes were then incubated in 20 μL of 0.4% Lucifer yellow-VS lithium salt in 60% L15 culture medium for 45 min at room temperature. Excess dye was removed by using a transfer pipette to drop in and remove the globes from 3 different Eppendorf tubes filled with 1000 μL of clean 60% L15. Rinsed globes were then fixed and prepared for imaging the same way as whole eyes for fluorescence immunohistochemistry (described above).

### Transmission Electron Microscopy (TEM)

Detailed methods for TEM tissue preparation are published elsewhere (21). Briefly, whole eyes were fixed in 4% paraformaldehyde + 1% glutaraldehyde in 0.1 M PB at 4°C for ≥ 24 hrs. Fixed eyes were then infiltrated with 2.3 M sucrose in 0.1 M PB for 1-4 hrs at 22°C with gentle shaking, embedded in OCT (ThermoFisher, Waltham, MA, USA), cryosectioned at 20 μm, and then thaw-mounted (one section per slide) onto gelatin-coated Fisherbrand Superfrost Plus slides. Optimally-oriented sections were washed with 0.1 M sodium cacodylate and then stained for 30 min with 1% osmium tetroxide. After staining, sections were dehydrated in increasing concentrations of anhydrous ethanol and then infiltrated with increasing concentrations of Eponate 12 resin (Ted Pella Inc., Redding, CA, USA) diluted with anhydrous EtOH. Once tissues were infiltrated with 100% resin, Beem^®^ capsules with the ends trimmed off were placed over the section on the slide and then the capsule was filled with resin and allowed to polymerize (16-24 hrs at 65°C). Ultrathin sections (silver-grey; 50-70 nm) were cut with a diamond knife and collected on 0.5% Formvar-coated nickel slot grids. Sections were stained with saturated aqueous uranyl acetate (12 min) and Venable and Coggeshall’s lead citrate (0.25%, 5 min). Imaging was performed with a Hitachi 7600 TEM at 80 kV.

### SDS-PAGE, Western blot, and dot blot

Single retinas from tadpoles aged > 45 days post-fertilization (minimum equatorial diameter ~1.6 mm) were isolated from the eye cup and solubilized in 50 μL of solubilizing buffer (1x PBS, 2.5% SDS, 5 mM Tris pH 6.8, 20% sucrose, bromophenol blue, 2 mM EDTA, 1 mM PMSF, 4% β-mercaptoethanol). Protein samples (12 μL) were separated with a 10% SDS-PAGE resolving gel using the Laemmeli discontinuous buffer system and then transferred to a PVDF transfer membrane (Immobilon-FL, Merck KGaA, Darmstadt, Germany) using a Biorad wet transfer apparatus. Blots were blocked for 30 min (1% skim milk in 1x PBS) and then probed with anti-N cdhr-21 (~99 kDa) or anti-N prom1 (~95 kDa) overnight. Blots were then incubated with IRDye800CW- or IRDye700CW-conjugated secondary antibodies (1:10,000; Rockland, Gilbertsville, PA) for 3 hrs and analyzed on a LI-COR Odyssey imager (Li-Cor, Lincoln, NE, USA).

### Electroretinography

Electroretinograms (ERGs) were recorded as previously described using electrodes connected to a model 1800 AC amplifier and head stage (AM Systems, Sequim, WA) and an Espion Ganzfeld stimulator (ColorDome) and recording unit (Diagnosys LLC, Lowell, MA) (22). Corneal recordings were made from a silver wire electrode set in a glass micropipette filled with 0.1x MMR and mounted into a micromanipulator. The combined reference and ground were a modified gold EEG electrode glued into a 60 mm petri dish. Tadpoles at Niewkoop and Faber stages 52-54 were anesthetized by exposure to 0.03% tricaine for 2 min and then mounted on their right side on the reference electrode using 2% low-melting point agarose infused with 0.01% tricaine. Scotopic ERGs were recorded in animals that had been dark-adapted overnight and prepared under dim red light; recorded stimuli were the average of 5 trials. Photopic ERGs were recorded in animals that had been exposed to a normal light cycle and prepared under regular lab lighting (350-500 lux); recorded stimuli were the average of 10 trials. All ERGs were recorded from the left eye, which was then fixed and processed for histology.

### Experimental Design & Statistical Analysis

Tadpole sex was not determined and phenotypic expression of mutations did not differ according to sex in frogs, so data from both sexes were pooled. The generation and number of animals used for each set of experiments are indicated in the results and in figure captions. Statistical analysis and graphing was performed using GraphPad Prism (V.6-8; San Diego, CA, USA). Western blot band density was quantified using FIJI (V 1.52p, Bethesda, Maryland, USA) and then unpaired, two-tailed t-tests were used to analyse differences between wildtype and mutant eyes. ERG waveforms were visualized in Excel (Microsoft, Redmond, WA) and analyzed by measuring the A-wave and B-wave peak amplitudes, plotting them, and then fitting the resulting curves using non-linear regression analysis. Genotype effects, light intensity effects, and interaction effects were analysed by two-way ANOVA with Sidak’s post hoc test. Photopic latency in *cdhr1*-null animals was analyzed by measuring the difference between peaks (*cdhr1*-null mutant minus wildtype) and then comparing the mean differences of all groups using a one-way ANOVA with Tukey post hoc test. Micrographs were processed using Adobe Photoshop (Creative Cloud 2019) and FIJI; any nonlinear adjustments in signal intensity are reported in the figure captions. Osmium peppering artifacts that obscured the underlying OS structure were digitally removed from some of the TEM micrographs. A preliminary report of some of our findings was presented previously in abstract form (Carr B, et al. IOVS 2019; 60 ARVO E-Abstract).

## RESULTS

CRISPR/Cas9-mediated gene editing is effective for *prom1* and *cdhr1* in *X. laevis* embryos. The *prom1* gene identified on xenbase.org (xelaev18034674mg, XB-GENE-6460662, NCBI Gene ID: 100316925 (*prom1.L*) corresponds to the *X. laevis* prominin-3 sequence, and is not expected to be expressed in high levels in the retina (23). The *prom1* sequence used in the experiments reported here – xelaev18005149m, NCBI Gene ID: 100316924 (*proml-1*) – was found by utilizing the xenbase.org BLAST database. It corresponds to the *X. laevis* prominin-1 sequence reported by Han and Papermaster and is therefore expected to have high retinal expression. There is likely no functional *prom1* S chromosome gene, as only 6/27 exons were identified in mRNA, protein, and EST databases; regardless, we did design one of the sgRNAs tested to target both the L and S chromosomes. Three *prom1* sgRNAs – targeting exon 1 (L), 12 (L & S), or 21 (L) – were tested and all resulted in successful editing. Sanger chromatograms for embryos with successful CRISPR/Cas9 editing were degraded near the predicted cut site, representing the occurrence of random indels due to nonhomologous end joining (NHEJ). The exon 1-targeting sgRNA was chosen for subsequent experiments because it had the highest editing efficiency (64% of eGFP+ embryos) and the resultant phenotype from all sgRNAs was the same. Three *cdhr1* guides – targeting exon 1 (L & S), 7 (L & S), or 8 (L & S) – were tested and only the exon 7-targeting sgRNA was successful in editing both the L (88% of eGFP+ embryos) and the S (32% of eGFP+ embryos) chromosomes. Reduction of prom1 and cdhr1 protein immunoreactivity was verified with Western blot and immunohistochemistry (*prom1*-null F0 & F1 and *cdhr1*-null F3, n = 10-12; Fig. 1). Genotypes for animals used in subsequent experiments were as follows: *prom1*-null F0: various random indels; *prom1*-null F1: L: homozygous 5 bp deletion, homozygous 6 bp deletion, or heterozygous 5 bp deletion + 6 bp deletion; *cdhr1*-null F2-3: L: homozygous 27 bp deletion, homozygous 10 bp deletion, or heterozygous 27 bp deletion + 10 bp deletion and S: homozygous 4 bp deletion;*prom1* + *cdhr1-null*: F0 various random indels.

**Figure 1.**
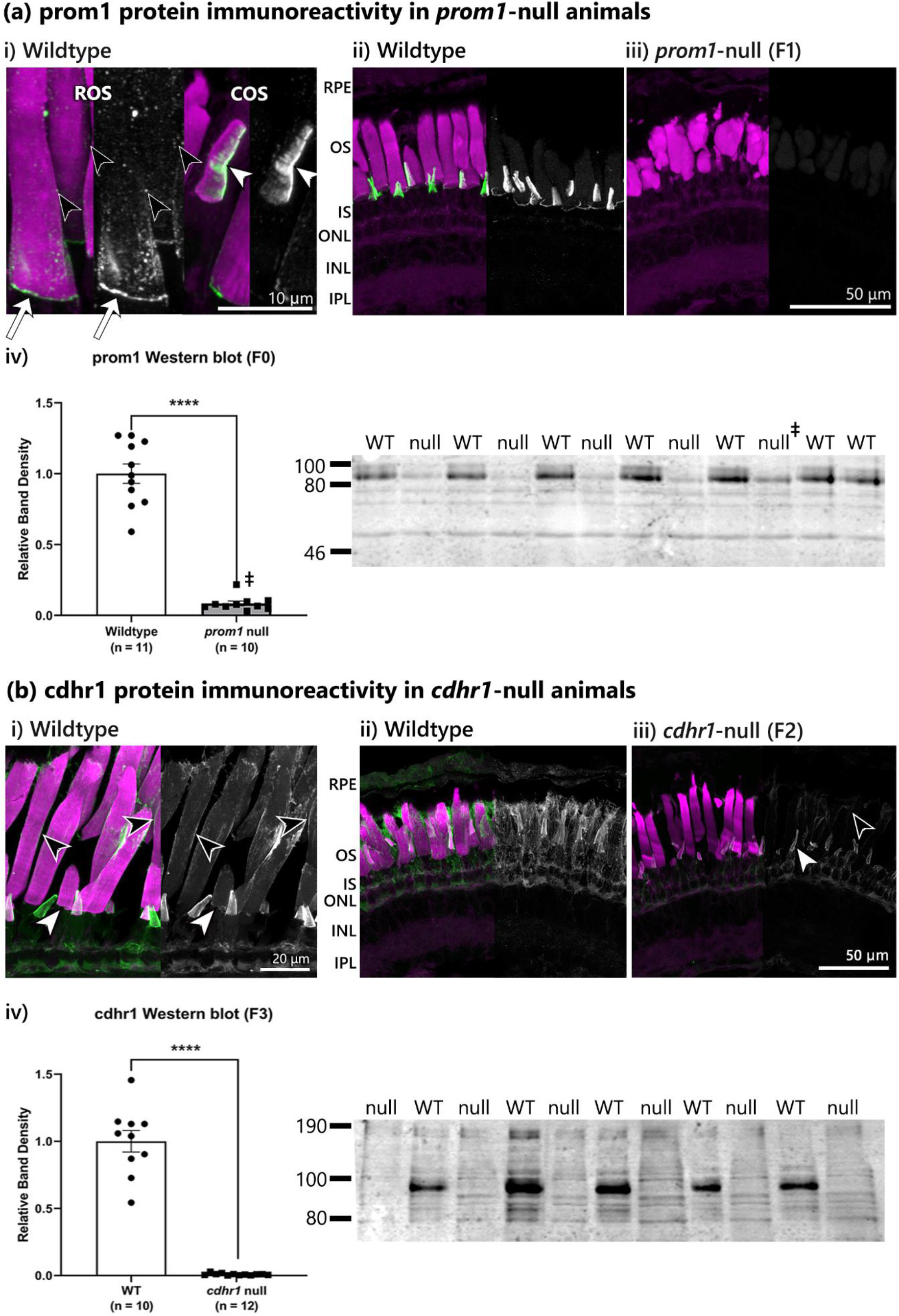
Comparison of wildtype animal immunoreactivity for prom1 and cdhr1 proteins to *prom1*-null animals (a) and *cdhr1*-null animals (b). (a i) In wildtype animals, prom1 is localized to the base of the ROS (white arrow), small puncta on the outside of the ROS (black arrowheads), and along one edge of the COS (white arrowheads). (a ii,iii) Prom1 immunoreactivity is lost in *prom1*-null retinas and ROS/COS are dysmorphic (F1, n = 14). (a iv) Western blot for *prom1*-null F0 animals demonstrates a significant reduction in prom1 protein immunoreactivity compared to wildtype (n = 10-11). One animal showed successful editing by Sanger sequencing but had a smaller reduction in prom1 immunoreactivity than the others (ǂ). (b i) In wildtype animals, cdhr1 protein is localized to a band at the base of the ROS (white arrowhead) and within the ROS plasma membrane (black arrowhead). (b ii,iii) ROS cdhr1 immunoreactivity is lost in *cdhr1*-null animals, but signal remains in the cone outer and inner segments and in presumptive RPE microvilli (F2, n = 11). (b iv) Western blot for *cdhr1*-null animals shows a complete reduction in cdhr1 protein immunoreactivity compared to wildtype in the band size that corresponds to the cdhr1 protein ~95 kDa (n = 10-12). Channels: magenta = WGA, green/white = prom1 (a) or cdhr1 (b). *Abbreviations*: WT = wildtype.

### Loss of *prom1* results in dysmorphic rods and severely dysmorphic cones

We found that prom1 protein is expressed in *X. laevis* retina in the basal ROS and in the COS opposite the disc rim protein peripherin-2 (prph-2), similar to results reported previously (10). We also found prom1-positive puncta scattered throughout the plasma membrane at the surface of the ROS, but not inside of the ROS (Fig. 1a, black arrowheads). CRISPR-mediated *prom1* gene knockdown significantly reduced retinal prom1 immunoreactivity in Western blot and immunohistochemistry, and resulted in dysmorphic photoreceptor OS (F0-F1, n = 10-26; Figs. 1, 2). In contrast to the uniform cylindrical wildtype ROS, *prom1*-null ROS were shortened and amorphous, with diameters changing markedly over short distances; these constrictions and bulges were comprised of overgrown or oddly grown OS disc membranes that were bent, folded, and formed circular whorls. There were also instances of overgrown loops of membranes that continued upwards alongside the ROS. The presence of prph-2 and lack of Lucifer yellow staining throughout the ROS indicates that although the ROS disc membranes are overgrown, they develop rims and do not remain open to the extracellular space (F0, n = 26, Fig. 2a,b). COS in *prom1*-null mutants were elongated and fragmented, and were often closely associated with the ROS plasma membranes – appearing to adhere to the ROS in fragmented puncta or wrap around the ROS in cone opsin positive tendril-like structures or sheets (F0 & F1, n = 14-26, Fig. 2c, asterisks & white arrowheads). There was occasional mislocalization of cone opsin to the inner segment plasma membrane in F0 animals, which varied greatly from a single cone inner segment to all of the cone inner segments in a section (15/42 specimens examined, 14-42 dpf; Fig. 2c). Cone opsin inner segment localization was most commonly correlated with complete destruction of the COS, although not always. Cone opsin mislocalization to the inner segment was not found in F1 animals with dysmorphic photoreceptors from two different genetic backgrounds (n = 10). There was no observed mislocalization of cdhr1 to the inner segment (F0 & F1, n = 14-26, Fig. 2a). Between 14 dpf to 6 weeks post-fertilization, dysmorphic ROS continued to grow to near adult length, but retained the dysmorphic structure of constrictions and bulges throughout the ROS; there was still no inner segment localization of any of the retinal proteins investigated. As animals aged, there was an increase in the appearance of small deposits that stained heavily with Hoescht and were also autofluorescent; thus, they are likely of cellular origin such as accumulations of cellular debris or possibly dying RPE cells (Fig. 2a, white arrowheads). These deposits did not occur in wildtype animals.

**Figure 2.**
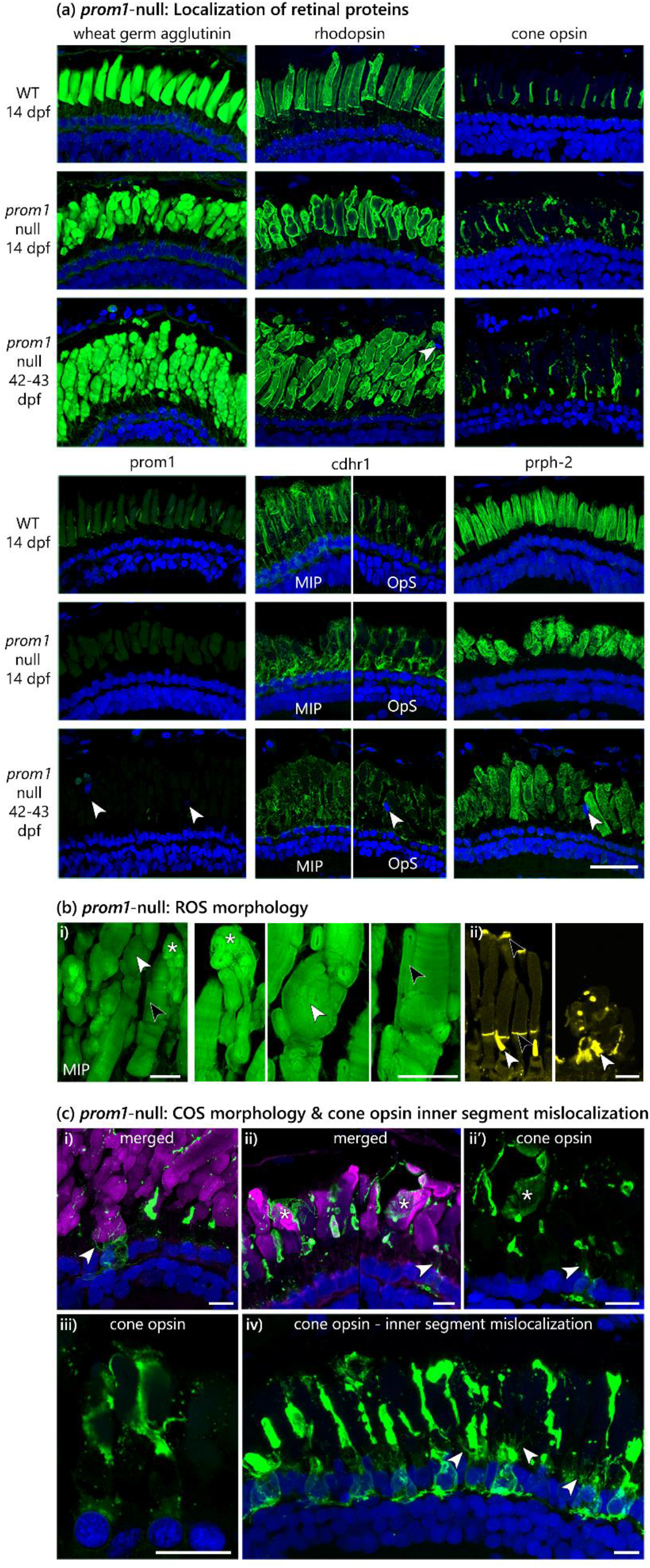
(a) Immunolabelling of various photoreceptor outer segment proteins in wildtype (14 dpf, top, n = 18) and F0 *prom1*-null (14 dpf, middle, n = 26; 42-43 dpf, bottom, n = 26) retinas. There is not mislocalization of any protein surveyed into the inner segment of the photoreceptors. As animals age, there is an increase in Hoechst-stained deposits in the outer segment layer (white arrowheads). *Channels*: green = protein of interest, blue = Hoechst. Scale bar = 50 μm. (b) *Prom1*-null ROS morphology. (b i) Structures of interest are: folded strings of membranes (white asterisk), large and small membrane whorls (white arrowhead), and overgrown folded OS membrane that is oriented vertically along the outside of the ROS (black arrowhead). (b ii) Lucifer yellow staining verified that the overgrown and dysmorphic ROS discs of *prom1*-null mutants are not open to the extracellular space (left = WT, right = prom1-null). COS are indicated by the white arrowheads and nascent ROS discs that are open to the extracellular space are indicated by black arrowheads. *Channels*: green = WGA, yellow = Lucifer yellow. Scale bar = 10 μm. (c) *Prom1*-null COS morphology. (c i-iv) Cone opsin positive membranes are fragmented and appear to be supported by neighbouring ROS. (c i, ii, iv) Tendrils of cone opsin-positive membrane are often seen wrapped around the base of adjacent ROS (white arrowheads) or draped over ROS in sheets (ii, white asterisks). (c iv) Cone opsin mislocalization to the inner segment occurs, but only in a small subset of animals (ca. 15% of retinas observed (F0), n = 41). *Channels*: green = cone opsin, blue = Hoechst, magenta = WGA. The green channel (cone opsin) was adjusted non-linearly in images c i, ii’, and iv to show the tendrils and sheets of cone opsin positive membrane and cone opsin inner segment localization with greater intensity. Scale bar = 10 μm. *Abbreviations*: COS = cone outer segment, dpf = days post-fertilization, MIP = maximum intensity projection, OpS = optical section (centre of stack), prph-2 = peripherin-2, ROS = rod outer segment, WT = wildtype.

When *prom1*-null retinas were examined using TEM, we found that ROS disc structural elements such as membrane laminations and hairpin rims were present and normal looking, even though the disc membranes themselves were highly disorganized. *Prom1*-null ROS disc membranes are convoluted with adjacent layers that run in different directions, but they remain contained by the plasma membrane. Overall, local disc structure was well-preserved, but higher-order disc organization was dramatically disrupted. Frequently observed features were turns, folds, strings of membrane, and whorls, and a there were also a few instances of penetration of disc membrane into the inner segment. In some ROS, the basal discs did not form hairpins; instead, they made a 90 degree turn from horizontal to the vertical direction, and then continued upwards along the ROS (F0, n = 3, Fig. 3a). In contrast, we found that COS disc membranes were very difficult to visualize with TEM due to their tortuous morphology and fragmentation of the membranes as observed using light microscopy. Almost no recognizable COS structures were observed by TEM other than occasional loops of disc membrane in close proximity to the COS. Most commonly, masses of completely disordered thin membranes above the inner segment were seen (F0, n = 3, Fig. 3b).

**Figure 3.**
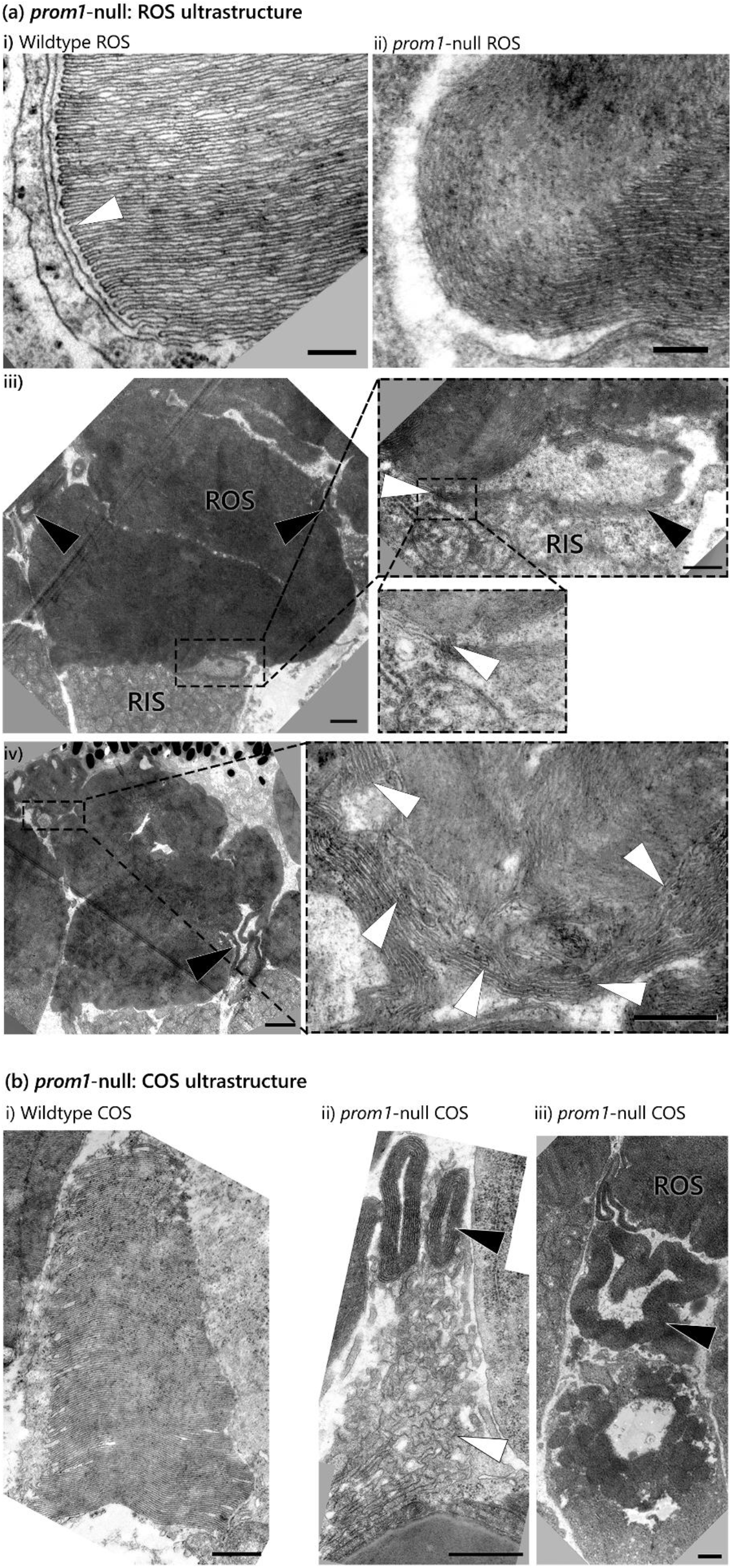
Transmission electron micrographs demonstrating the principal changes in ultrastructure of *prom1*-null mutants compared to wildtype controls. (a i) Wildtype ROS ultrastructure is highly ordered and consists of stacked OS membrane discs with properly formed hairpins (white arrowhead). Prom1-null ROS lack hairpins in the proper areas (a ii) and have a severely convoluted structure where the disc membranes appear to be bent over, folded, or exist in thin tracts (a iii, black arrowheads). There are some instances of disc membranes invaginating into the rod inner segment (a iii inset; black arrowhead = membrane discs, white arrowhead = hairpins). There are commonly hairpins at the top of the ROS (iv inset, white arrowheads = hairpins, black arrowhead = thin tract of disc membrane). (b i) Wildtype COS consist of cone-shaped ordered stacks of lamellae. (b ii, iii) Common features of prom1-null COS are loops of disc membrane that appear unattached to the CIS (ii-iii, black arrowhead), and the presence of thin, convoluted membranes above the CIS (ii, white arrowhead). Scale bar = 800 nm. *Abbreviations*: COS = cone outer segment, RIS = rod inner segment, ROS = rod outer segment.

### Loss of *prom1* results in impaired cone function

The impact of *prom1*-null mutations on the scotopic ERG in 6 week old tadpoles was minimal. At the lowest light intensities, the scotopic A-wave amplitude was reduced compared to wildtype controls (a 75-78% reduction in response amplitude at 2.5-25 cd/m^2^; simple main genotype effect, F (1, 78) = 13.13, p = 0.0005), but the difference between *prom1*-null and wildtype A-wave amplitudes decreased as light intensities increased (a 44-53% reduction at 250-750 cd/m^2^, no difference at 1250-2500 cd/m^2^). *Prom1*-null scotopic B-wave amplitudes were not significantly different from wildtype controls, and no change in the shape of the scotopic ERG waveform was observed (n = 7; Fig. 4).

**Figure 4.**
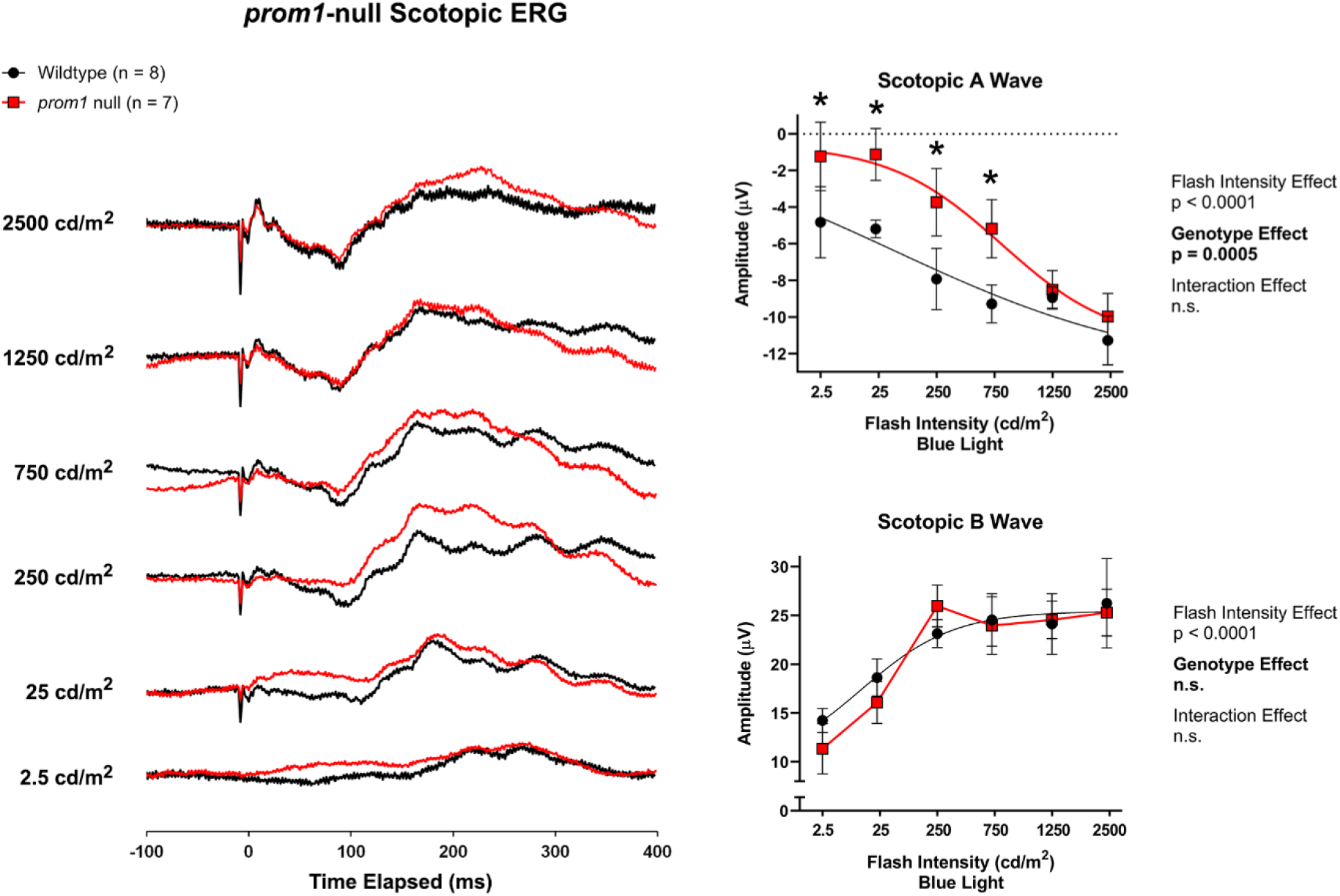
Averaged scotopic single-flash recordings from wildtype and *prom1*-null F0 animals (n = 7-8). Waterfall plots (left) and transformed linear regression curves (right) were used to visualize and compare wildtype and *prom1*-null ERG waveforms and A- and B-wave amplitudes. Data analysis utilized a Two-Way ANOVA with Sidak’s post hoc test. Data are plotted as mean ± SEM. *Statistics*: p < 0.05 *.

The impact of *prom1*-null mutation on the photopic ERG of 6 week old tadpoles was significant. *Prom1*-null photopic A- and B-wave amplitudes were significantly reduced at higher light intensities relative to wildtype controls. For the photopic A-wave (interaction effect, F (5, 72) = 3.435, p = 0.0077), there was no difference at 0.25-7.5 cd/m^2^ and a 30-50% reduction at 25 and 75 cd/m^2^; (simple main genotype effect, F (1, 72) = 4.686, p = 0.0337). For the photopic B-wave (interaction effect F (5, 72) = 6.290, p < 0.0001), there was no difference from 0.25-2.5 cd/m^2^, a 46% reduction at 7.5 cd/m^2^, and a 64-66% reduction at 25-75 cd/m^2^ (simple main genotype effect, F (1, 72) = 34.29, p < 0.0001). *Prom1*-null cone response to 5 Hz photopic flicker was also reduced relative to wildtype at higher light intensities (interaction effect, F (5, 72) = 4.179, p = 0.0022). There was no difference from 0.25-0.75 cd/m^2^, a 58% reduction at 2.5-7.5 cd/m^2^, and a 50-54% reduction at 25-75 cd/m^2^ (simple main genotype effect, F (1, 72) = 34.84, p < 0.001) (F0, n = 7, Fig. 5).

**Figure 5.**
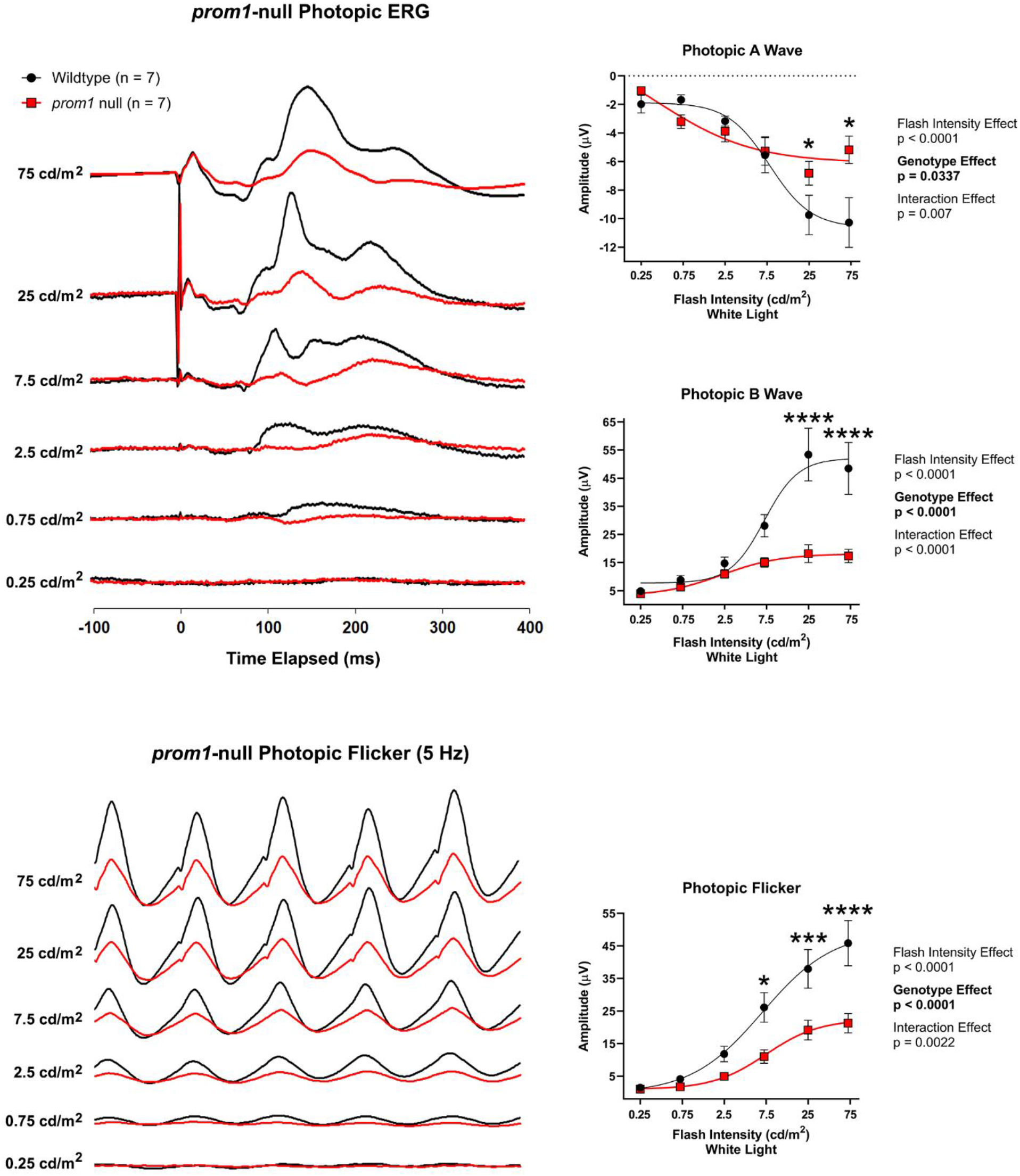
Averaged photopic single-flash and 5 Hz photopic flicker recordings from wildtype and *prom1*-null F0 animals (n = 7). Waterfall plots (left) and transformed linear regression curves (right) were used to visualize and compare wildtype and *prom1*-null A-wave, B-wave, and flicker responses. Data analysis utilized a Two-Way ANOVA with Sidak’s post hoc test. Data are plotted as mean ± SEM. *Statistics*: p < 0.05 *, p < 0.001 ***, p < 0.0001 ****.

### Loss of *cdhr1* results in OS disc orientation and growth defects

We determined that *X. laevis* cdhr1 protein is expressed at the basal ROS and in the ROS plasma membrane. *Cdhr1*-null animals lost cdhr1 immunoreactivity in the ROS, but some signal remained in the COS and possible RPE microvilli (F2-3, n = 7-11, Fig. 1b). This immunoreactivity is likely not cdhr1-specific; Western blots demonstrated loss of one strong band of the expected size in *cdhr1*-null animals, but also showed the presence of several non-specific bands (F3, n = 12, Fig. 1b). *Cdhr1*-null mutants did not have severely dysmorphic photoreceptor OS, and differences between mutants and wildtype animals were not easily detectable by regular confocal microscopy. Possible subtle changes could be a narrowing and lengthening of the ROS and some COS that were shortened or collapsed at the tips. The expression and localization of rhodopsin, cone opsin, prom1, and prph-2 were all normal. Using super-resolution microscopy, we observed that some rod photoreceptors (~20 %) had areas of membrane overgrowth that extended from the basal ROS upwards, alongside the outside of the regularly-ordered disc membranes and incisures. There was also visible “pock-marking” or holes in some of the basal ROS. There was no penetration of Lucifer yellow into the ROS, however, indicating that although there is abnormal membrane growth, the disc membranes are sealed off from the extracellular space. (F2-3, n = 5-11, Fig. 6b).

**Figure 6.**
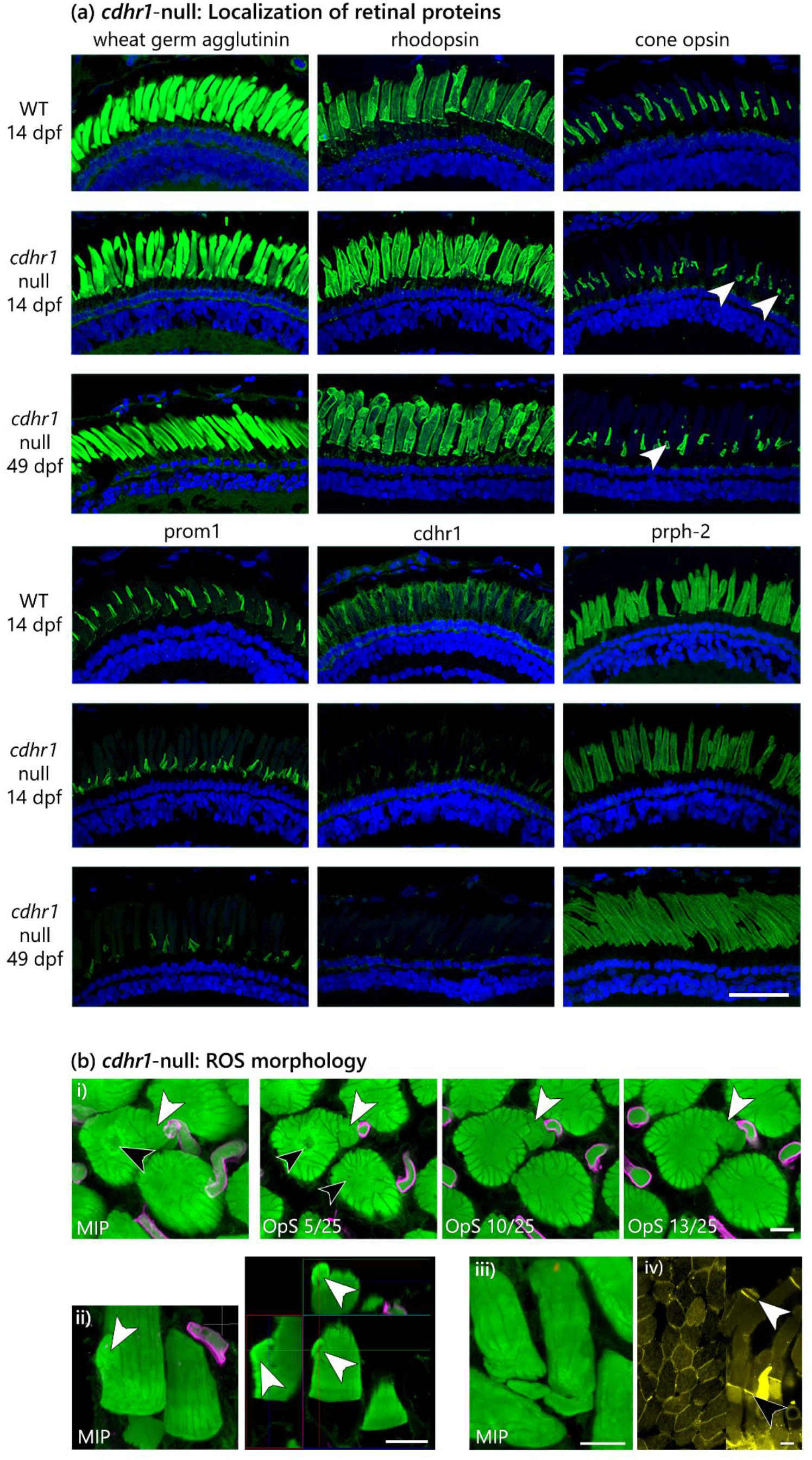
Immunolabelling of photoreceptor outer segment proteins in wildtype (14 dpf, top; n = 10) and F3 *cdhr1*-null (14 dpf, middle, n = 11; 42-43 dpf, bottom, n = 7) retinas. *Channels*: green = protein of interest, blue = Hoechst. Scale bar = 50 μm. (b) Three different examples of overgrown membranes visualized by super-resolution microscopy in *cdhr1*-null ROS. (b i) A maximum intensity projection (left) and 4 optical sections (right) demonstrating the structure of a membrane overgrowth in a basal coronal section of ROS. Features of interest are the overgrown membrane (white arrowheads) and the holes or “pock-marking” at the base of the OS looking up from the inner segment (MIP & OpS 5/25, black arrowheads). (b ii) A side-view of overgrown disc membranes which appear to be comprised of a large overgrowth that folds back onto itself. (b iii) A long “tail” of overgrown disc membrane that extends from, and then loops under, the basal ROS. (b iv) Lucifer yellow staining of *cdhr1*-null ROS in the coronal (left) and sagittal (right) orientations; nascent discs are normally open to the extracellular space (black arrowheads), as are discs near the tip of the ROS (white arrowheads), but there is no abnormal Lucifer yellow dye penetration into the ROS. *Channels*: green = WGA, magenta = cone opsin, yellow = Lucifer yellow. Scale bars = 5 μm. *Abbreviations*: dpf = days post-fertilization, prph-2 = peripherin-2, WT = wildtype.

Ultrastructural analysis by TEM confirmed disc membrane orientation and growth defects in the ROS. The principal defect observed was that some disc membranes (~30%) were oriented vertically within the ROS plasma membrane; these defects occurred both as long, thin sections of vertically-oriented disc membranes and shorter “stacked” thicker sections of vertical membranes comprised of short pieces of disc membrane and many rim structures. Rim structures and “bubbles” of membrane – which likely correspond to the “pock-marking” seen in the super-resolution images – were almost always present at the point at which disc orientation was altered. Horizontally- and vertically-oriented ROS membrane discs had normal lamination and the discs remained tightly packed into the ROS plasma membrane. COS discs appeared relatively free of defects, although some COS (~10%) had a “frayed” appearance, in which disc lamella were not uniformly registered, suggesting over- or under-growth of the disc membranes at somewhat regularly-spaced intervals (F0-F1, 14-49 dpf, n = 5; Fig. 7).

**Figure 7.**
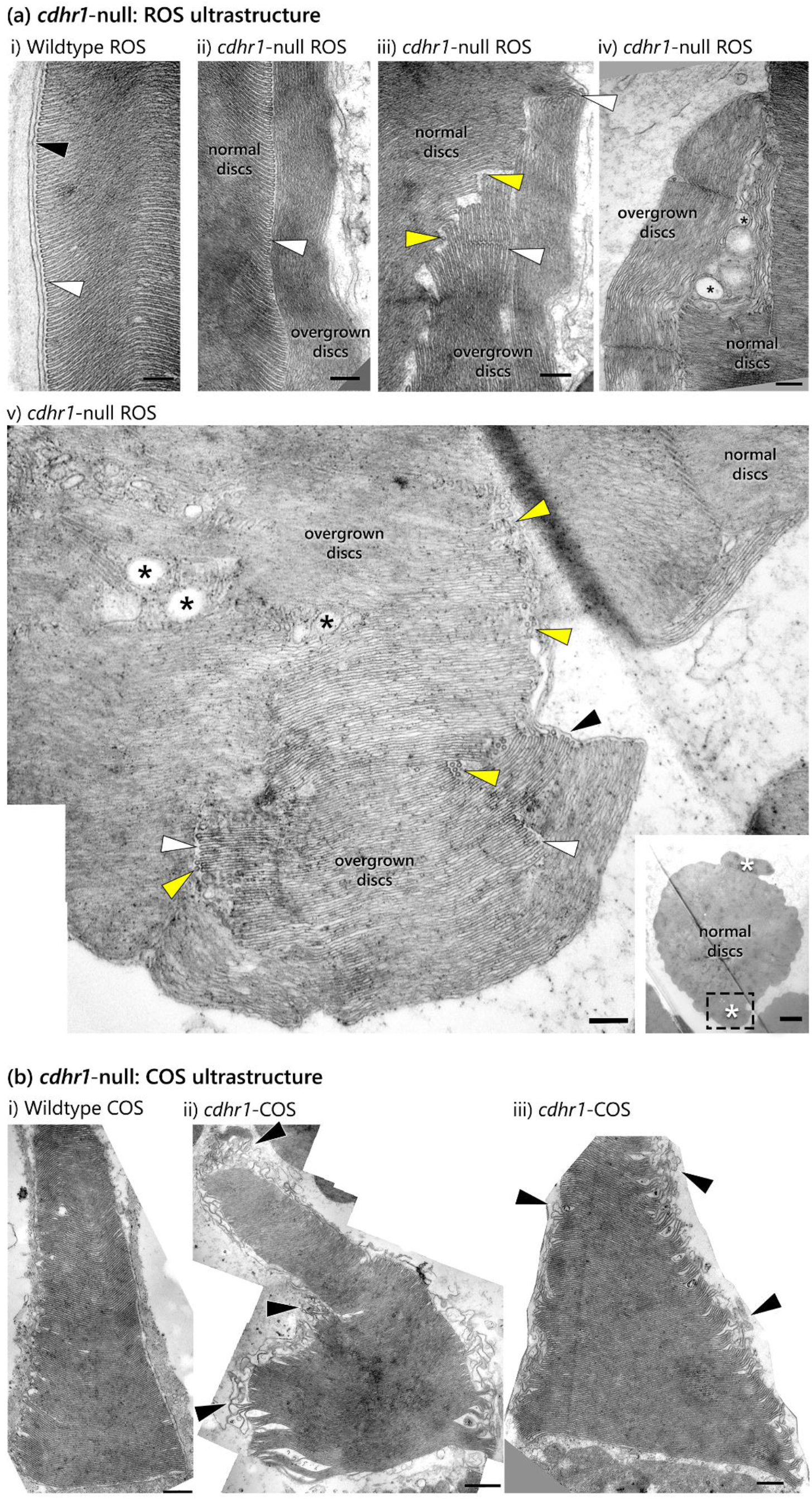
Transmission electron micrographs demonstrating the changes in ultrastructure of *cdhr1*-null mutants compared to wildtype controls. (a i) Wildtype ROS have organized structure with horizontal disc membranes and neatly-aligned hairpins. (a ii-iv) The principal feature of *cdhr1*-null ROS are overgrown discs that are oriented vertically instead of horizontally within the ROS plasma membrane. (a iii-v) Areas of overgrowth are commonly associated with large (asterisks) or small (yellow arrowheads) bubbles of membranes around the area where disc orientation changes occur. (a i-iii, v) Hairpins (white arrowheads) are present where disc membranes are overgrown and these areas of disc overgrowth or disorganization appear to be contained within the plasma membrane (v, black arrowhead). Overgrowth of disc membrane is easily seen when the ROS is in the coronal orientation (v, inset, white asterisks). (b i) Wildtype COS ultrastructure is also organized, with neatly-stacked and aligned disc membrane lamellae. (b ii-iii) Although the ultrastructure of COS is mostly retained, there are some examples of COS where the disc membranes are over- or under-grown, so that there is a loss of registration of disc membranes, giving the COS a frayed appearance. F0 & F3, n = 5; Scale bar = 200 nm except (v, inset), where the scale bar = 2 μm. *Abbreviations*: COS = cone outer segment, ROS = rod outer segment.

### Loss of cdhr1 may affect photoreceptor signalling kinetics

The shape and scale of the scotopic ERG from six week old *cdhr1*-null tadpoles was not significantly different from wildtype animals in regards to the A- or B-wave amplitude, but the B-wave tended to return to baseline more quickly in *cdhr1*-null animals (F3, n = 8, Fig. 8). There was no statistically significant effect on photopic ERG amplitudes, although there was a trend towards a slightly larger B-wave response and a small increase in latency for B-wave onset; this latency was not statistically significant for any condition other than 5 Hz flicker at 25 cd/m^2^ (one-way ANOVA, F (5, 24) = 6.625, P = 0.0005) (F3, n = 10, Fig. 8).

**Figure 8.**
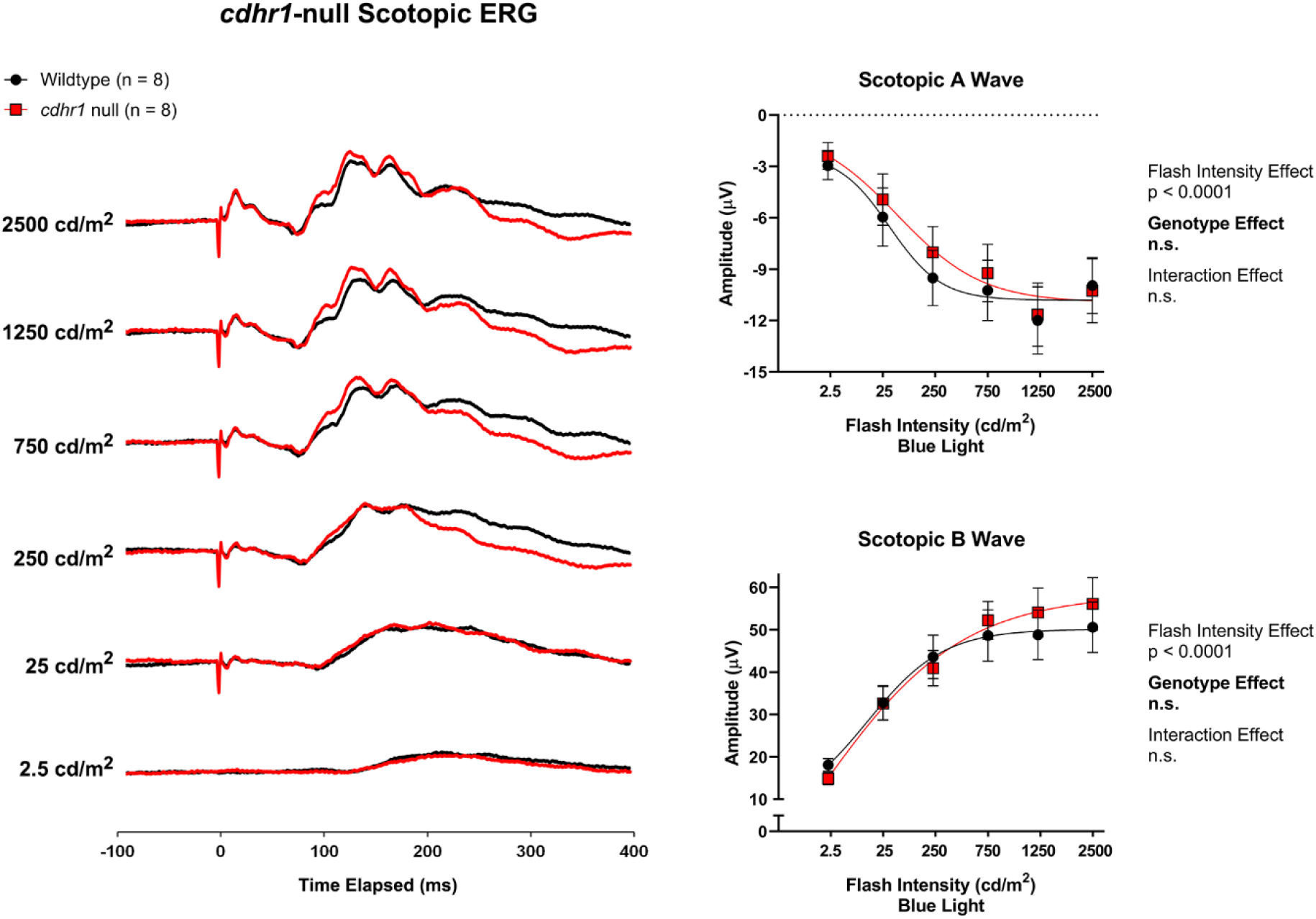
Averaged scotopic single-flash recordings from wildtype and *cdhr1*-null F3 animals (n = 8). Waterfall plots (left) and transformed linear regression curves (right) were used to visualize and compare wildtype and *cdhr1*-null A-wave and B-wave responses. Data analysis utilized a Two-Way ANOVA with Sidak’s post hoc test. Data are plotted as mean ± SEM.

**Figure 9.**
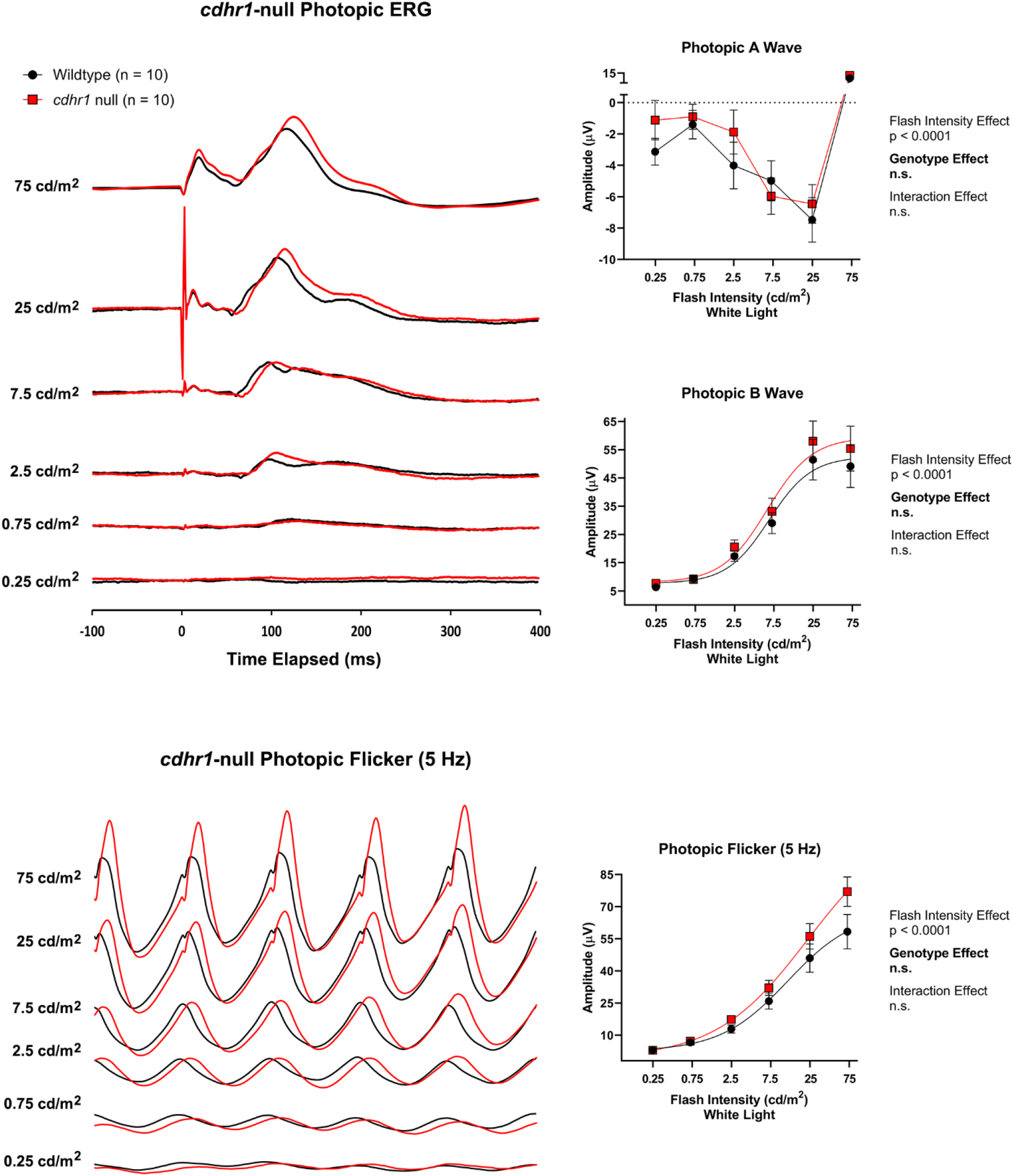
Averaged photopic single-flash and 5 Hz flicker recordings from wildtype and *cdhr1*-null F3 animals (n = 10). Waterfall plots (left) and transformed linear regression curves (right) used to visualize and compare wildtype and *cdhr1*-null A-wave, B-wave, and flicker responses. The large positive A-wave values at 75 cd/m^2^ is likely an artefact introduced by the large early receptor potential response measured by the electrode used in this experiment. Data analysis utilized a two-way ANOVA with Sidak’s post hoc test. Data are plotted as mean ± SEM.

Combination of *prom1* + *cdhr1* knockdown does not result in a more severe phenotype or more severe functional impairment than *prom1* knockdown alone.

Photoreceptors with both *prom1* and *cdhr1* gene knockdown were not significantly more dysmorphic or prone to degeneration than *prom1*-null animals. The effects on OS structure and protein localization were not distinguishable from *prom1*-null retinas for both light microscopy and TEM (F0, n = 3-14; Fig. 10, Fig. 11). Older double-null animals also had the small Hoechst-stained autofluorescent deposits in the OS layer (Fig. 10).

**Figure 10.**
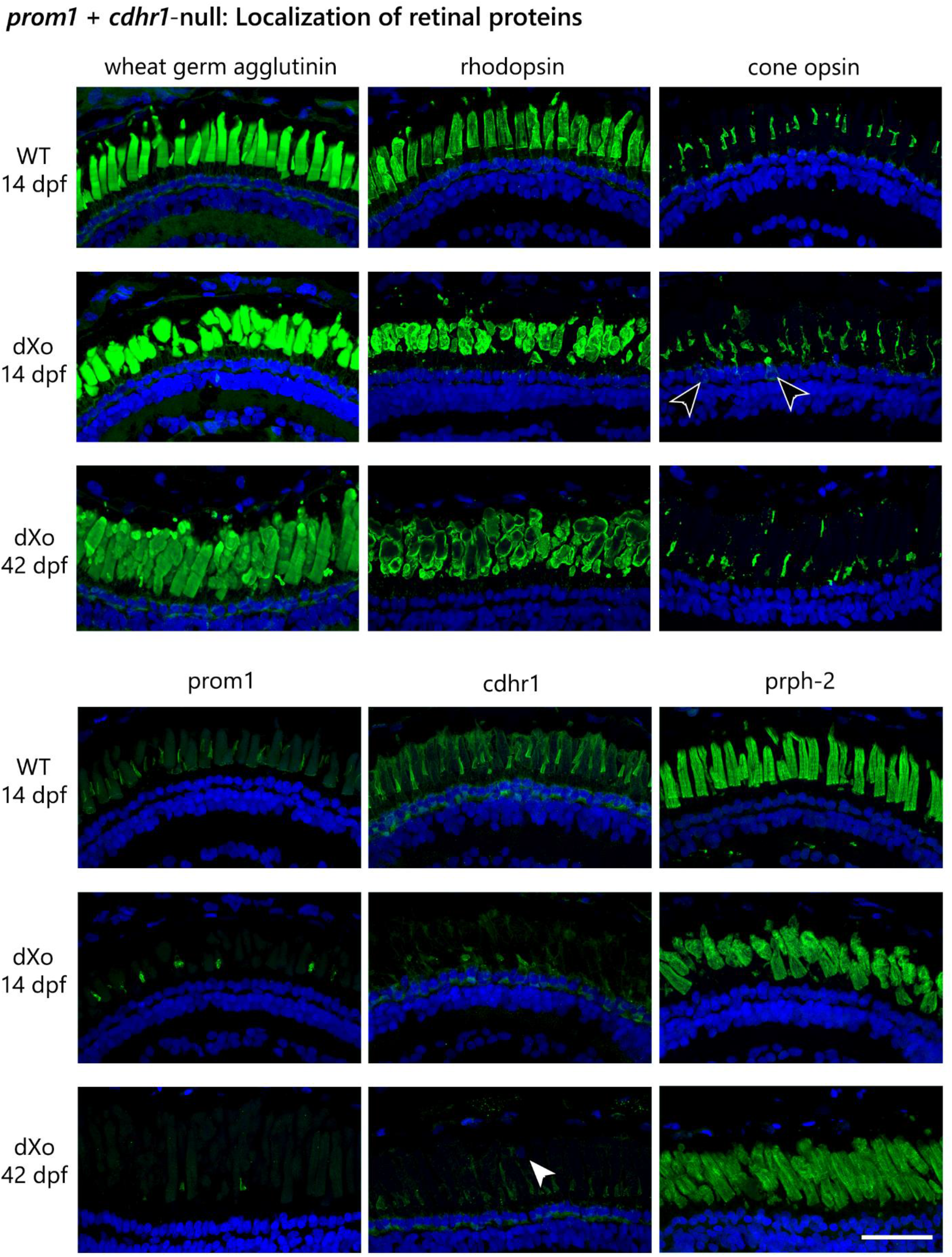
Immunolabelling of various photoreceptor outer segment proteins in wildtype (14 dpf, top; n = 11) and F0 *prom1 + cdhr1*-null (14 dpf, middle, n = 14; 42-43 dpf, bottom, n = 7) retinas. Similar to *prom1*-null animals, there are occasional instances of mislocalization of cone opsin to the inner segment (black arrowheads) and an increase in condensed nuclei in the outer segment layer in older animals (white arrowhead). *Channels*: green = protein of interest, blue = Hoechst. Scale bar = 50 μm. *Abbreviations*: dpf = days post-fertilization, dXo = double-null (*prom1* + *cdhr1*-null), prph-2 = peripherin-2, WT = wildtype.

**Figure 11.**
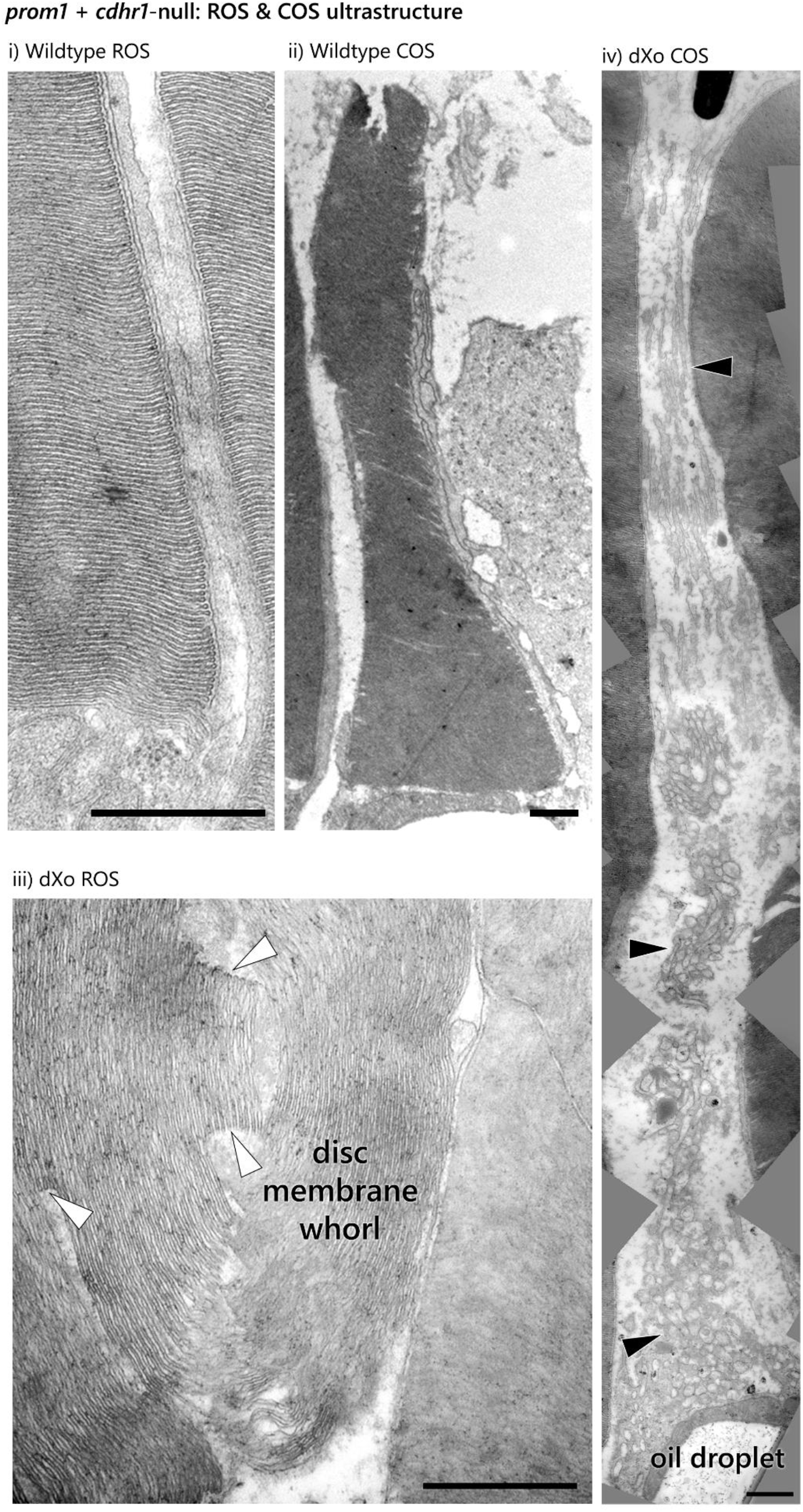
The ultrastructure of ROS and COS in *prom1* + *cdhr1*-null mutant animals. OS defects are the same as those that were observed in prom1-null mutants; ROS are overgrown, convoluted, and membrane discs are comprised of folds, whorls, and stacks of membrane that contain hairpins (iii, white arrowheads). COS are similarly difficult to visualize, but are comprised of loose, convoluted and looped thin membranes (iv, black arrowheads). Scale bar = 800 nm. *Abbreviations*: COS = cone outer segment, dXo = double-null (*prom1* + *cdhr1*-null), ROS = rod outer segment.

The impact of *prom1* + *cdhr1*-null mutations on photoreceptor function was similar to that of *prom1*-null mutations. Double-null mutants were less sensitive to lower and moderate intensity scotopic stimuli but the difference decreased at higher intensity stimuli (scotopic A wave: no difference in response amplitude from baseline at 2.5-25 cd/m^2^, a 76% reduction at 250 cd/m^2^, no difference at 750-2500 cd/m^2^; simple main genotype effect, F (1, 35) = 6.511, p = 0.0152). There was no significant difference in scotopic B-wave amplitude or change in the shape of the scotopic ERG waveform (n = 3-5; Fig. 12).

**Figure 12.**
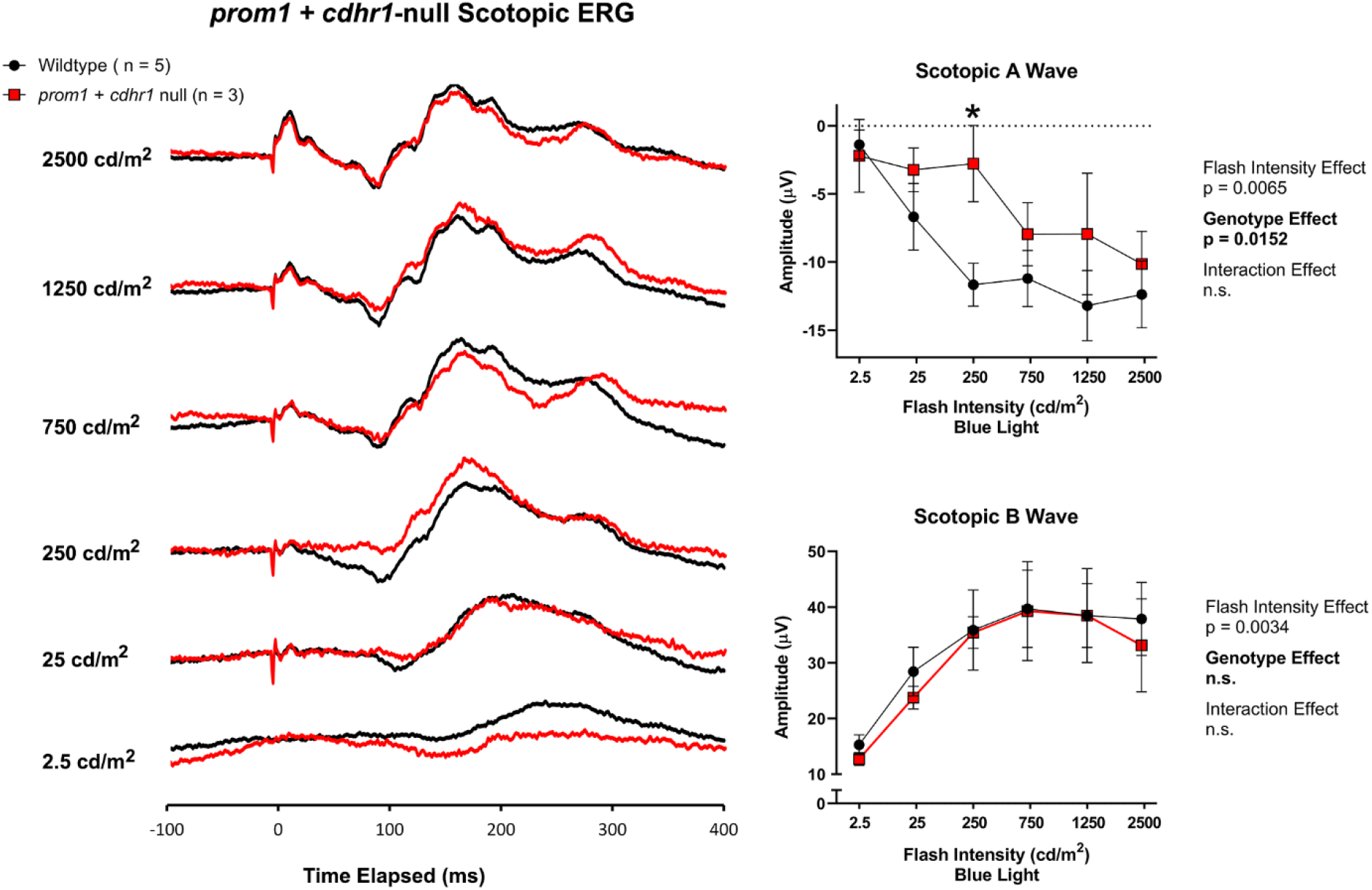
Averaged scotopic single-flash recordings from wildtype and *prom1* + *cdhr1*-null F1 animals (n = 3-5). Waterfall plots (left) and transformed linear regression curves (right) were used to visualize and compare wildtype and *prom1* + *cdhr1*-null A-wave and B-wave responses. Data analysis utilized a Two-Way ANOVA with Sidak’s post hoc test. Data are plotted as mean ± SEM. *Statistics*: p < 0.05 *.

Cone function of *prom1 + cdhr1*-null mutants was impaired compared to WT animals, and the effects were similar to those of *prom1*-null mutants; the photopic A-wave response to increasing white light intensities was reduced. The A-wave (no difference from 0.25-7.5 cd/m^2^, a 42% reduction at 25 cd/m^2^, and a 77% reduction at 75 cd/m^2^; simple main genotype effect, F (1, 42) = 7.861, p = 0.0076) and the B-wave response curves were flattened (interaction effect, F (5, 42) = 12.93, p < 0.0001; no difference from 0.25-2.5 cd/m^2^, a 44% reduction at 7.5 cd/m^2^, a 56% reduction at 25 cd/m^2^, and a 54% reduction at 75 cd/m^2^; simple main genotype effect, F (1, 42) = 62.69, p < 0.0001). The response to photopic flicker (5 Hz) of *prom1* + *cdhr1*-null animals was also reduced (no difference from 0.25-2.5 cd/m^2^, a 50% reduction at 7.5 cd/m^2^, a 37% reduction at 25 cd/m^2^, and a 33% reduction at 75 cd/m^2^ (interaction effect, F (5, 42) = 3.641, p = 0.0080; simple main genotype effect, F (1, 42) = 27.67, p < 0.0001) (F0, n = 4-5, Fig. 13).

**Figure 13.**
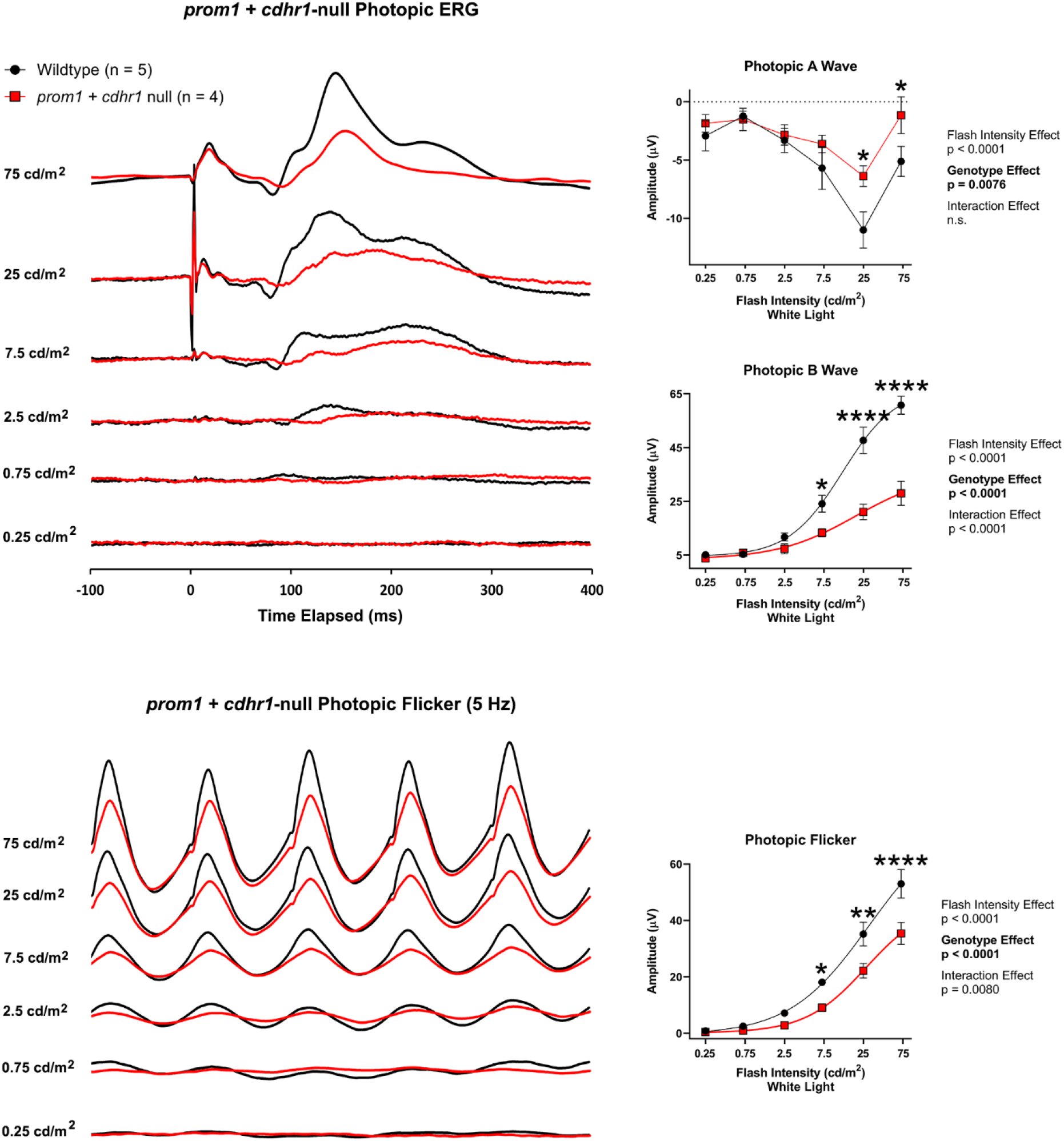
Averaged photopic single-flash and 5 Hz flicker recordings from wildtype and *prom1* + *cdhr1*-null F1 animals (n = 4-5). Waterfall plots (left) and transformed linear regression curves (right) were used to visualize and compare wildtype and *prom1* + *cdhr1*-null A-wave, B-wave, and flicker responses. Data analysis utilized a two-way ANOVA with Sidak’s post hoc test. Data are plotted as mean ± SEM. *Statistics*: p < 0.05 *, p < 0.01 **, p < 0.0001 ****.

## DISCUSSION

The central finding of this study is that neither *prom1* nor *cdhr1* are necessary for photoreceptor outer segment disc membrane evagination, disc fusion, or the maintenance of the spacing of disc membrane lamellae. Our results suggest that *prom1* and *cdhr1* have distinct roles in regulating different aspects of nascent outer segment disc membrane size and organization. Prom1 may regulate disc size, by aligning and reinforcing interactions between the leading edges, while cdhr1 may regulate disc membrane organization by helping to keep the discs horizontal before fusion occurs. Different roles for prom1 and cdhr1 in OS disc morphogenesis is supported by the observation that phenotypes caused by *prom1*-null mutations are significantly more severe than those resulting from *cdhr1*-null mutations and there is no mislocalization to the inner segment of prom1 in *cdhr1*-null animals or cdhr1 in *prom1*-null animals. A secondary important finding of this study is that the retinas of *prom1-, cdhr1-*, and double-null *X. laevis* do not degenerate quickly. The maintenance and growth of ROS, the lack of any OS protein mislocalization to the inner segment, the preservation of photoreceptor function, and the increasing appearance of autofluorescent deposits in the OS layer of older *prom1*-null animals suggests that retinal degeneration caused by these mutations may be due to secondary toxic retinal effects – e.g., RPE toxicity or accumulation of cellular waste products – and not due to direct effects of these mutations on OS structure or the improper trafficking of OS proteins. This is an important finding, as it may not be critical to prevent the dysmorphic photoreceptors caused by prom1-null mutations to preserve vision; therapies could instead be targeted to prevent the secondary events that ultimately cause cell death.

The effects of *prom1*-null mutations on photoreceptor structure are severe. The presence of convoluted overgrown membranes indicates that prom1 likely plays a role in regulating the size of OS membrane discs, possibly by controlling the amount of membrane that is added before disc fusion occurs or by aligning and reinforcing interactions between the leading edges of the discs as they elongate. Nascent disc evagination and disc fusion still occurs in *prom1*-null mutants, as evidenced by the lack of photoreceptor death or degeneration of the OS and by the presence of hairpins and the lack of Lucifer yellow staining within the ROS. Recent studies have suggested that prom1 may be involved in cytoskeletal remodelling, specifically by interacting with phosphoinositide 3-kinase and the Arp2/3 complex (16), and a conditional knockdown of the Arp2/3 complex in mice ROS results in abnormal OS structure with a similar phenotype to that of *prom1-null X. laevis*; OS membranes form knob-like protrusions made up of whorls of overgrown disc membranes (24). Overgrowth of disc membranes also occurs when eyecups are treated with cytochalasin D, a mycotoxin that inhibits actin polymerization (25). At the molecular level, prom1 has been reported to associate with actin and to regulate membrane localization and retention of cholesterol (18, 26). Cholesterol reinforces positive membrane curvature, such as the leading edges of evaginating discs, and loss of membrane rigidity in the leading edges of the COS lamella could explain the elongated and fragmented appearance of *prom1*-null *X. laevis* cones; unlike ROS, the COS do not have the extra structural support of full disc fusion or a surrounding plasma membrane.

The survival of dysmorphic photoreceptors in *X. laevis* supports that *prom1* is not required for biosynthesis of OS discs or trafficking of opsins to the OS. There is no mislocalization of key OS proteins such as rod opsin, cone opsins, or prph-2 to the inner segment, and *prom1*-null *X. laevis* lack the severe retinal degeneration frequently associated with defects in ciliary or trafficking components (5). Instead, the earlier and severe retinal degeneration reported in mice and zebrafish could be due to the shorter lifespans of these animals compared to *X. laevis* (1.5-3.5 yrs vs. 15-30 yrs), and the retinal degeneration reported could be compounded by age-related retinal degenerative components instead of being directly caused by the loss of *prom1*. Cone opsin mislocalization occurred only in a small subset of *prom1*-null *X. laevis*, unlike previous reports in *prom1^-/-^* and *prom1^R373C^* mice (11, 18), and our results indicate that this mislocalization is usually the result of complete destruction of the COS. The lack of early and severe retinal degeneration in *prom1*-null *X. laevis* supports the hypothesis that disruption of OS disc morphogenesis is not the cause of photoreceptor death, but that an indirect secondary effect could be responsible instead. In support of this, there are increasing numbers of small, heavily Hoechst-stained autofluorescent deposits in the OS layer of 6 week old *prom1*-null *X. laevis*. An increase in lipofuscin-like deposits in *prom1^R373C^* mice has also been reported (18). Clinically, some *prom1* mutant retinal diseases resemble Stargardt disease (6, 7, 27), which is also caused by secondary toxic effects – i.e., the build-up of bisretinoid A2PE (lipofuscins) due to the lack of ABCA4 kills the RPE, leading to subsequent photoreceptor death (28, 29). It should be noted, however, that *prom1*-null and *prom1^R373C^* patients lack the retinal hyperfluorescence associated with ABCA4-associated Stargardt disease.

The *cdhr1*-null phenotype in *X. laevis* is less severe than the effects of *prom1*-null mutations and the *cdhr1*-null phenotype reported in mouse. OS disorganization is limited to changes in disc membrane orientation, poor disc stacking, and occasional membrane whorls due to oversized disc membranes. There is no mislocalization of OS proteins such as rhodopsin, cone opsins, prph-2, or prom1 and the lack of Lucifer yellow dye penetration into the overgrown ROS membranes indicates that they are sealed off from the extracellular space. The subtle changes in ERG response suggest that all components required for phototransduction are present and functional. *X. laevis* photoreceptors appear largely unaffected by the loss of *cdhr1*, even though mice were reported to experience up to 50% loss of outer nuclear layer density by 6 months of age (8). This difference in reported photoreceptor death between species could be the result of differences in cdhr1 protein cleavage and localization in the ROS. In mice, cdhr1 is localized only to the basal ROS, where it is cleaved into a soluble N-terminus and a membrane-embedded C-terminus (19). It was hypothesized that this cleavage represents an irreversible step in rod OS morphogenesis, such as during membrane fusion when nascent open discs are closed off and enclosed within the ROS plasma membrane. *Xenopus* discs appear to require neither cdhr1 nor cdhr1 cleavage for membrane fusion however – N-terminal immunoreactivity is present in the basal ROS as well as throughout the ROS plasma membrane in wildtype animals and hairpins are detectable using TEM in *cdhr1*-null mutants – suggesting that this process is not integral to fusion of the leading edges. It has also been suggested that cdhr1 may act as a tether between the leading edge of nascent ROS discs and the inner segment, and that it could guide OS disc growth until the disc has reached the correct size, after which the tether is severed (2). Our data support this hypothesis over the cleavage hypothesis. *X. laevis* have calyceal processes made up of F-actin fibres that form a cage-like structure around the base of the rod and cone OS, which mice lack (30). Possibly, the presence of calyceal processes could lessen the impact of the loss of cdhr1 if its function is to tether nascent discs in alignment, as they could provide additional structural support and guidance for nascent disc membranes as they elongate.

Our study does not support the existence of a prom1-cdhr1 protein complex that performs a single, shared role in OS disc morphogenesis as previously hypothesized (18). If this were true, then it would be expected that *prom1*-null and *cdhr1*-null mutations should affect OS morphology similarly. Instead, the *prom1*-null phenotype is significantly more severe than *cdhr1*-null phenotype and there is no mislocalization of prom1 in *cdhr1*-null animals or cdhr1 in *prom1*-null animals. These proteins may still interact, but their function is not dependent on the presence of the other protein. Double-null mutants do not have a significantly different phenotype than *prom1*-null mutants, which provides further evidence against a genetic or protein interaction. If a relationship existed, we would expect that double-null animals would have either a significantly more severe phenotype (synergy) or a mitigated phenotype, which may occur when gene products operate in series within the same pathway (31).

In summary, the results reported in this study have provided significant new insights into the function of prom1 and cdhr1 proteins in photoreceptor outer segment morphogenesis and the pathogenesis of *prom1*-null mutations in human disease. Our data support a role for prom1 in the regulation of nascent disc size and structural support for the OS and a role of cdhr1 in disc membrane tethering and organization. Our study also shows definitively that these proteins are not required for outer segment disc evagination, the formation of hairpins, or disc fusion. Furthermore, we are the first to report that *prom1*-null mutations may cause retinal degeneration by secondary effects instead of direct effects on photoreceptor OS morphogenesis. This new insight could lead to a paradigm shift in the development of therapies for human patients, as it may not be necessary to rebuild the photoreceptors to preserve vision; therapies could instead be targeted to preventing the secondary events that ultimately cause cell death. This is an exciting subject of future investigation.

## Conflict of Interest Statement

The authors declare that they have no conflicts of interest with the contents of this article.

## Acknowledgements

Research funded by the Canadian Institutes of Health Research (OLM; PJT-155937, PJT-156072), the Natural Sciences and Engineering Research Council of Canada (OLM; RGPIN-2015-04326), the Edwina and Paul Heller Memorial Fund (OLM/BJC), and a Michael Smith Foundation for Health Research Research Trainee Award (BJC; 18367). Purchase of the Zeiss LSM 800 and LSM 880 with Airyscan funded by an infrastructure grant from the Canadian Foundation for Innovation (OLM). Transmission electron microscopy imaging supported by the University of British Columbia Bioimaging Facility and staff.

## ABBREVIATIONS

OS: outer segment
ROS: rod outer segment
COS: cone outer segment
prom1: prominin-1
cdhr1: cadherin-related family member 1 (photoreceptor cadherin)
ERG: electroretinography
RPE: retinal pigment epithelium
WT: wildtype
WGA: wheat germ agglutinin
sgRNA: single-guide RNA
TEM: transmission electron microscopy.

